# Induction of T-DNA amplification by retrotransposon-derived sequences

**DOI:** 10.1101/2023.03.05.531200

**Authors:** Lauren Dickinson, Wenxin Yuan, Chantal LeBlanc, Geoffrey Thomson, Siyuan Wang, Yannick Jacob

**Author notes:** These authors contributed equally to this manuscript (Lauren Dickinson, Wenxin Yuan).

## Abstract

Transformation via *Agrobacterium tumefaciens* (Agrobacterium) is the predominant method used to introduce exogenous DNA into plants. Transfer DNA (T-DNA) originating from Agrobacterium can be integrated as a single copy or in concatenated forms in plant genomes, but the mechanisms affecting final T-DNA structure remain unknown. In this study, we demonstrate that the inclusion of retrotransposon (RT)-derived sequences in T-DNA can increase transgene copy number by more than 50-fold in *Arabidopsis thaliana* (Arabidopsis). RT-mediated amplification of T-DNA results in large concatemers in the Arabidopsis genome, which are primarily induced by the long terminal repeats (LTRs) of RTs. T-DNA amplification is dependent on the activity of DNA repair proteins associated with theta-mediated end joining (TMEJ). Finally, we show that T-DNA amplification can increase the frequency of targeted mutagenesis and gene targeting. Overall, this work uncovers molecular determinants that modulate T-DNA copy number in Arabidopsis and demonstrates the utility of inducing T-DNA amplification for plant gene editing.

## Main Text

T-DNA integration into plant genomes via Agrobacterium-mediated transformation is used for a myriad of applications in plant biology, including the introduction of gene editing components and sequences conferring traits for crop improvement. In some instances, single copy insertions of T-DNA are desirable as they are expected to produce more consistent phenotypes between independent transgenic lines. However, it has long been recognized that multicopy T-DNA insertions in the form of concatemers are frequent outcomes of plant transformation^1–4^. A better understanding of these amplified T-DNA structures could potentially be advantageous for specific applications that would benefit from increased transgene copy number. For example, DNA repair templates required for gene targeting can be delivered via T-DNA, and *in vivo* levels of these templates have been shown to correlate positively with the frequency of homology-directed repair^5^. While much progress has been made in understanding the mechanisms responsible for chromosomal integration of T-DNA in plants, relatively little is known regarding how T-DNA copy number may be experimentally regulated.

As a strategy to increase T-DNA copy number in Arabidopsis, we explored the possibility that including retrotransposon (RT)-derived sequences within a T-DNA may induce transgene amplification. In plants, RTs and other repetitive sequences are preferentially amplified over protein-coding genes in the absence of the histone mark H3.1K27me1^6^, a process that depends on DNA repair^7^. In addition, RTs contain sequence features that can interfere with replication, transcription, and DNA repair, which can lead to locus-specific genomic amplification^8^. To assess if RT-derived DNA sequences increase T-DNA copy number, we designed a T-DNA vector based on the well-studied ONSEN family of long terminal repeat (LTR) RTs^9, 10^. We first replaced part of the *gag* and *pol* genes of an ONSEN RT (*At1g11265*) with DNA from the Arabidopsis *ACETOLACTATE SYNTHASE* (*ALS*) locus, with the aim of using this *ALS* fragment 1) to measure T-DNA copy number relative to the Arabidopsis genome and 2) to serve as a repair template in gene targeting assays (Fig. 1a). The resulting ONSEN RT was then subcloned into the T-DNA region of a binary vector modified from a previously described gene targeting study (see Methods section)^11^. Identical plasmids, except for the presence of ONSEN sequences surrounding *ALS*, were then used to transform Arabidopsis Col plants via Agrobacterium using the floral dip method (Fig. 1b)^12^. Genomic DNA was extracted from individual first-generation transformed (T1) plants and *ALS* quantification was performed by quantitative PCR (DNA-qPCR) using a primer set that can amplify both endogenous and T-DNA-associated *ALS* sequences. We found that T1 plants transformed with the plasmid containing ONSEN sequence (ONSEN RT) had, on average, higher *ALS* copy numbers compared to untransformed plants or T1 plants transformed with the control plasmid lacking ONSEN (No RT) (Fig. 1c). Individual T1 plants transformed with ONSEN RT showed wide variation in *ALS* levels, with copy number sometimes reaching 50-fold increase relative to Col. In contrast, T1 plants transformed with the plasmid lacking ONSEN rarely showed more than 5-fold increase over Col (Fig. 1c). To determine if other parts of the T-DNA were being amplified, we quantified the relative levels of the *pea3A* terminator (*pea3A(T)*) and the kanamycin- resistance *NPTII* cassette present at the 5’ and 3’ regions of the T-DNA, respectively (Fig. 1b). We detected similar increases in copy numbers for *ALS*, *pea3A(T)*, and the *NPTII* cassette in individual T1 plants transformed with ONSEN RT (Fig. 1d), indicating that the entire T-DNA, and not solely ONSEN-embedded *ALS*, was amplified.

**Figure 1.**
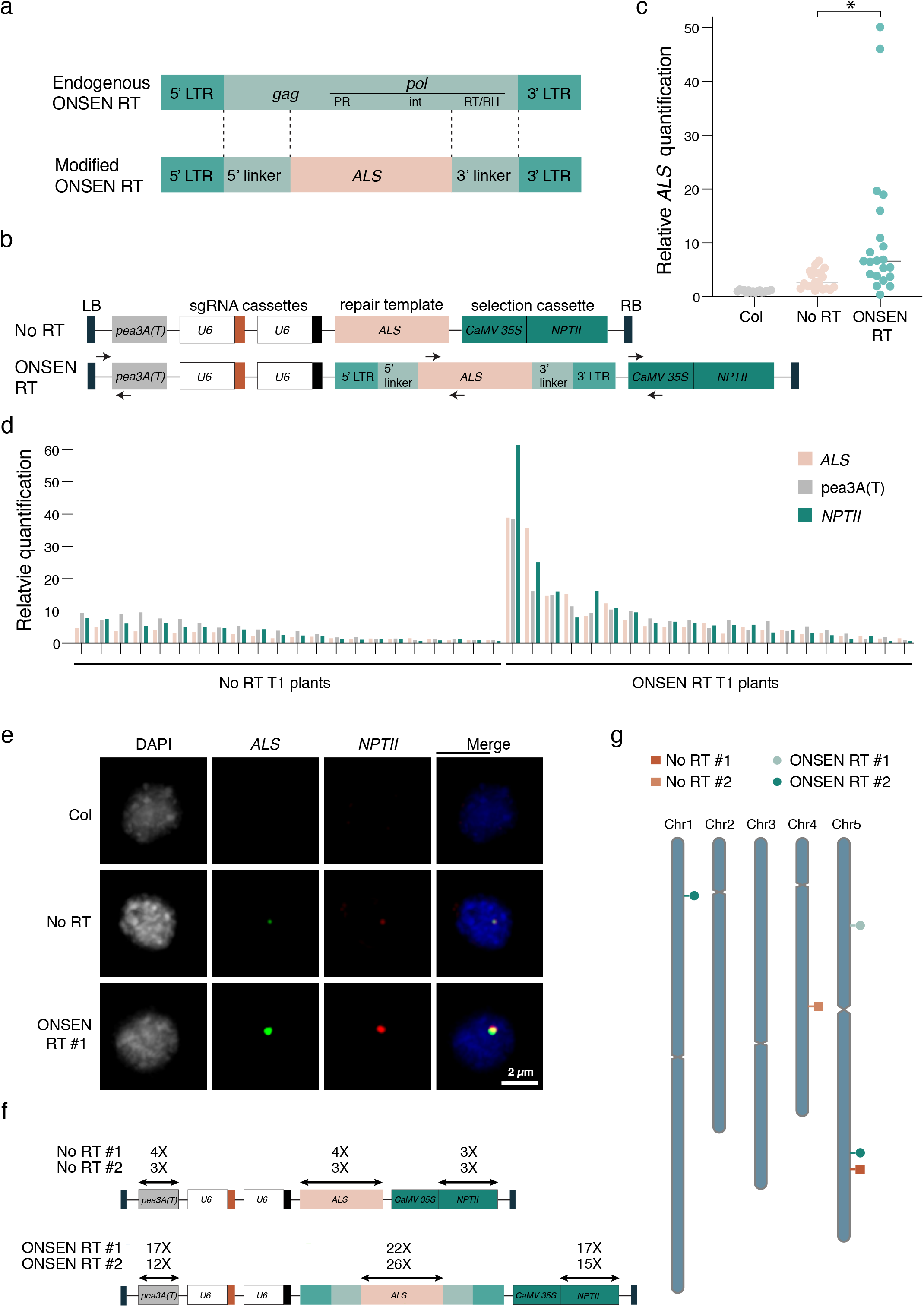
T-DNAs featuring retrotransposon sequences are amplified in Arabidopsis. **a.** Structure of the endogenous ONSEN RT and the modified RT used in the ONSEN RT-based vector. LTR: long terminal repeat; gag: gag-like protein; PR: protease; int: integrase; RT/RH: reverse transcriptase/RNase H. **b.** Schematic representation of the No RT (adapted from^11^) and ONSEN RT T-DNAs. Arrows indicate the primers used for real time quantification shown in panels c and d. LB: left border; RB: right border. **c.** DNA-qPCR of *ALS* in Col, No RT, and ONSEN RT T1 plants. Each dot represents an individual plant. Horizontal bars indicate the median. \**P* < 0.001 (Mann-Whitney *U* test). **d.** DNA-qPCR of *ALS*, *pea3A(T)* and *NPTII*. **e.** Fluorescence *in situ* hybridization in leaf nuclei using probes for *ALS* (green) and *NPTII* (red). Nuclei were stained with DAPI. **f.** Transgene copy number determined by whole genome sequencing analysis. **g.** T- DNA insertion sites in No RT and ONSEN RT T1 plants.

To characterize the structural organization of the ONSEN-derived T-DNA in the Arabidopsis genome, we performed DNA fluorescence *in situ* hybridization (FISH) experiments with probes designed to detect *ALS* and *NPTII*. In nuclei of T1 plants transformed with ONSEN RT and confirmed by DNA-qPCR to contain large numbers of T-DNA copies (>10-fold vs Col) (Extended Data Fig. 1a), we observed bright and overlapping signals for *ALS* and *NPTII* (Fig. 1e and Extended Data Fig. 1b). We typically detected 1-2 FISH signals per nucleus, which is in line with the average number of T-DNA integrations when performing Arabidopsis floral transformation^13–15^. By contrast, FISH signals in nuclei from Col or T1 plants transformed using the plasmid lacking ONSEN were either undetectable or much weaker (Fig. 1e and Extended Data Fig. 1b). The fact that overlapping and high-intensity FISH signals can be observed for *ALS* and *NPTII* strongly suggests that ONSEN-induced T-DNA amplification results from concatenation rather than an increase in T-DNA insertion sites in the Arabidopsis genome. To validate this, we sequenced the genome of a Col control and four T1 plants (two plants transformed with each type of T-DNA plasmid) characterized by different levels of T-DNA amplification as assessed by DNA-qPCR (Extended Data Fig. 2a). Analysis of the sequencing data confirmed the increase in copy number of the T-DNA and showed that these plants contained only one or two T-DNA insertion sites (Fig. 1f-g and Extended Data Fig. 2b), thus supporting that T-DNA amplification is due to increased copies of T-DNA via concatenation and not additional T-DNA insertion sites in the genome.

Next, we investigated the mechanism(s) involved in ONSEN-mediated T-DNA amplification. First, we assessed if T-DNA amplification could be conferred by sequences from RTs other than ONSEN. We tested four different Arabidopsis LTR RTs (two Copia-type and two Gypsy-type) by replacing part of their *gag-pol* sequence with the same *ALS* fragment present in ONSEN RT (Fig. 2a and Fig. 1a) and observed a similar effect of increased T-DNA copies compared to the ONSEN RT plasmid (Fig. 2b). We then investigated which region(s) of the ONSEN RT are involved in T-DNA amplification by generating a series of binary plasmids containing various deletions of the ONSEN RT plasmid (Fig. 2c). Our results show that the LTRs had the most impact on amplification (Fig. 2d). We hypothesized that the effect of the LTRs on T-DNA amplification is mainly caused by their repetitive nature, which we confirmed by assessing T-DNA levels induced by a binary plasmid with random repeated sequences (RR) of the same length and GC content as the ONSEN LTRs (Fig. 2e-f). Interestingly, not all DNA direct repeats within a T-DNA contribute to amplification, as removal of one of two identical sgRNA genes, which eliminates a repeated Arabidopsis U6-26 promoter sequence of 387 bp, had no effect on T-DNA copy number (Extended Data Fig. 3). Thus, multiple pairs of DNA repeats, and/or a linker sequence of defined length separating individual repeats, may be required to induce T-DNA amplification.

**Figure 2.**
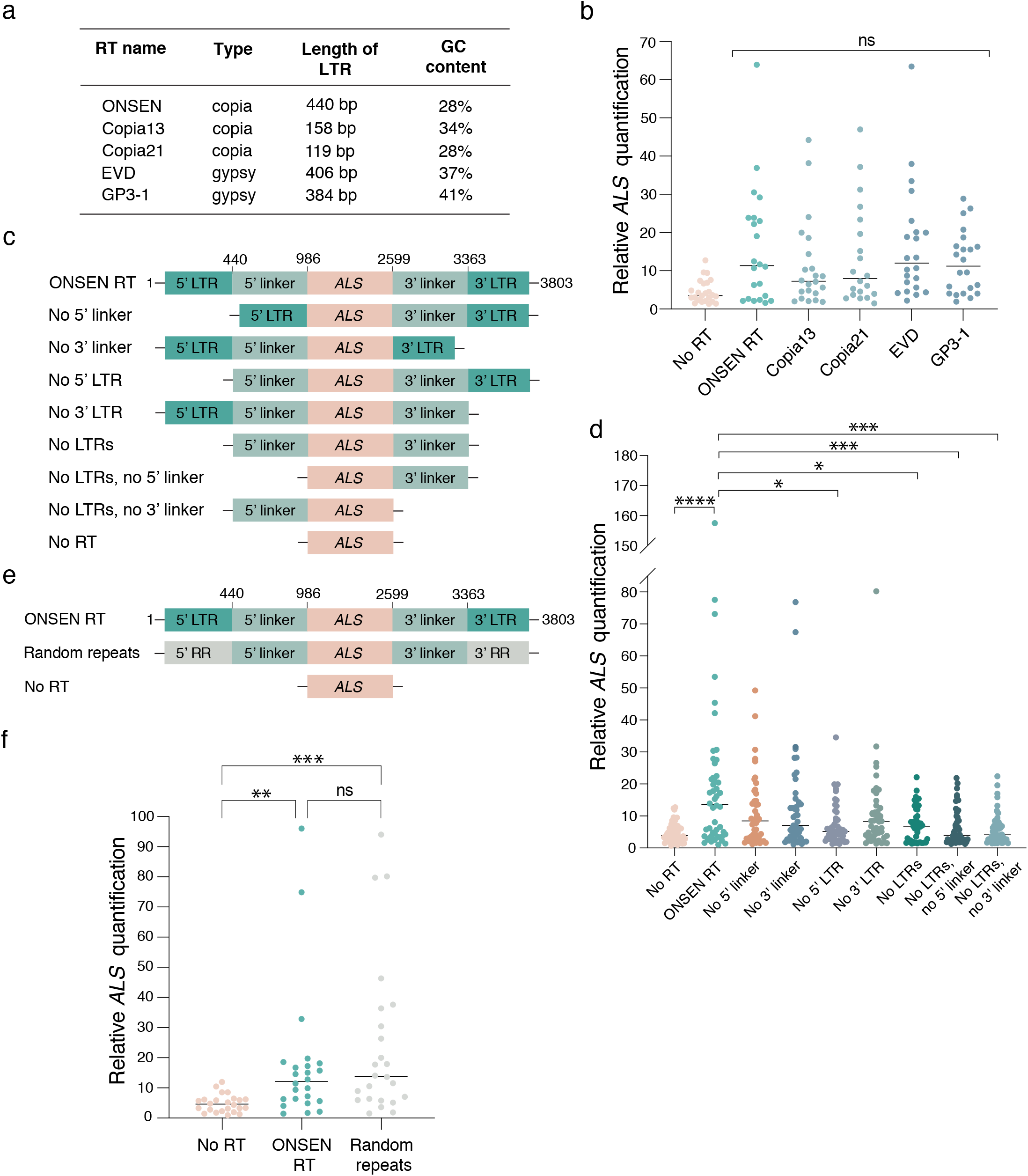
The repetitive nature of LTRs induces T-DNA amplification. **a**. Classification and characteristics of retrotransposons used to make RT-based T-DNAs. **b.** DNA-qPCR of *ALS* in plants transformed using various RT-based plasmids. Each dot represents an individual T1 plant. Horizontal bars indicate the median. ns = not significantly different (Kruskal-Wallis ANOVA followed by Dunn’s test). **c.** Schematic representations of ONSEN deletion constructs used in panel d. **d.** DNA-qPCR of *ALS* in plants transformed with ONSEN RT deletion constructs. Each dot represents an individual T1 plant. Horizontal bars indicate the median. \**P* < 0.05, \*\**P* < 0.01, \*\*\**P* < 0.001, \*\*\*\**P* < 0.0001 (Kruskal-Wallis ANOVA followed by Dunn’s test). **e.** Schematic representation of random repeat construct used in panel f. **f.** DNA-qPCR of *ALS* in plants transformed with ONSEN RT and random repeat constructs. Each dot represents an individual T1 plant. Horizontal bars indicate the median. \*\**P* < 0.01, \*\*\**P* < 0.001, ns = not significantly different (Kruskal-Wallis ANOVA followed by Dunn’s test).

T-DNA amplification may originate in Agrobacterium or arise before, during or after T-DNA integration in the plant genome. Our results indicate that T-DNA levels are similar in cultured Agrobacterium strains carrying plasmids with or without RT sequences (Fig. 3a), arguing that T-DNA amplification takes place within the plant. Furthermore, whole-genome sequencing analysis did not reveal copy number variations for the genomic regions flanking the T-DNA insertion sites (Fig. 3b), suggesting that amplification occurs at the time of, or before, T-DNA integration. In Arabidopsis, two DNA repair pathways contribute to T-DNA integration: TMEJ and non-homologous end joining (NHEJ)^16, 17^. TMEJ is an error-prone repair pathway responsible for inducing large (>5 kb) tandem genomic amplifications^18^, as observed in Arabidopsis mutants lacking the histone mark H3.1K27me1 (e.g., *atxr5 atxr*6 mutant^19, 20^) that are characterized by amplification of heterochromatin in a manner dependent on the replication fork-repair factor TONSOKU (TSK)^7^. To test if RT-mediated T-DNA amplification is caused by specific DNA repair pathways, we transformed our binary plasmids (with and without ONSEN) into several mutant backgrounds. First, we tested NHEJ by transforming *ku70*, *ku80* and *lig4* mutants, but we did not detect a significant effect on T-DNA amplification (Fig. 3c). To assess TMEJ, we could not directly verify the involvement of DNA polymerase theta (the main component of this repair pathway), as T-DNA integration is abolished in the absence of this protein^17^. Therefore, we assessed mutant backgrounds of TMEJ-associated RAD17 and MRE11^16, 21^, and observed strong suppression of T-DNA amplification (Fig. 3d). RAD17 and MRE11 also contribute to homologous recombination (HR)-mediated repair^22–24^, but, using a *rad51* mutant background, we found that HR is not involved in the induction of T-DNA amplification (Fig. 3e). In support of the involvement of TMEJ in T-DNA amplification, we identified mutational signatures consistent with this repair pathway (e.g., microhomology-associated deletions and small filler sequence insertions^18^) in sequencing reads spanning T-DNA copy junctions (Extended Data Fig. 4). Further investigations of DNA repair proteins revealed that the damage-induced kinase ATR, but not ATM, is required to increase T-DNA copy number (Fig. 3f). Finally, in contrast to heterochromatin amplification in the absence of H3.1K27me1, T-DNA amplification is not affected by mutations in *ATXR5*/*ATXR6* or *TSK*, thus suggesting that the amplification mechanism is not dependent on DNA replication (Extended Data Fig. 5a). In accordance, T-DNA levels are relatively constant in tissues of T1 plants separated by large number of replication cycles (Extended Data Fig. 5b). In sum, our results support a key role for TMEJ in inducing T-DNA amplification before or during T-DNA integration.

**Figure 3.**
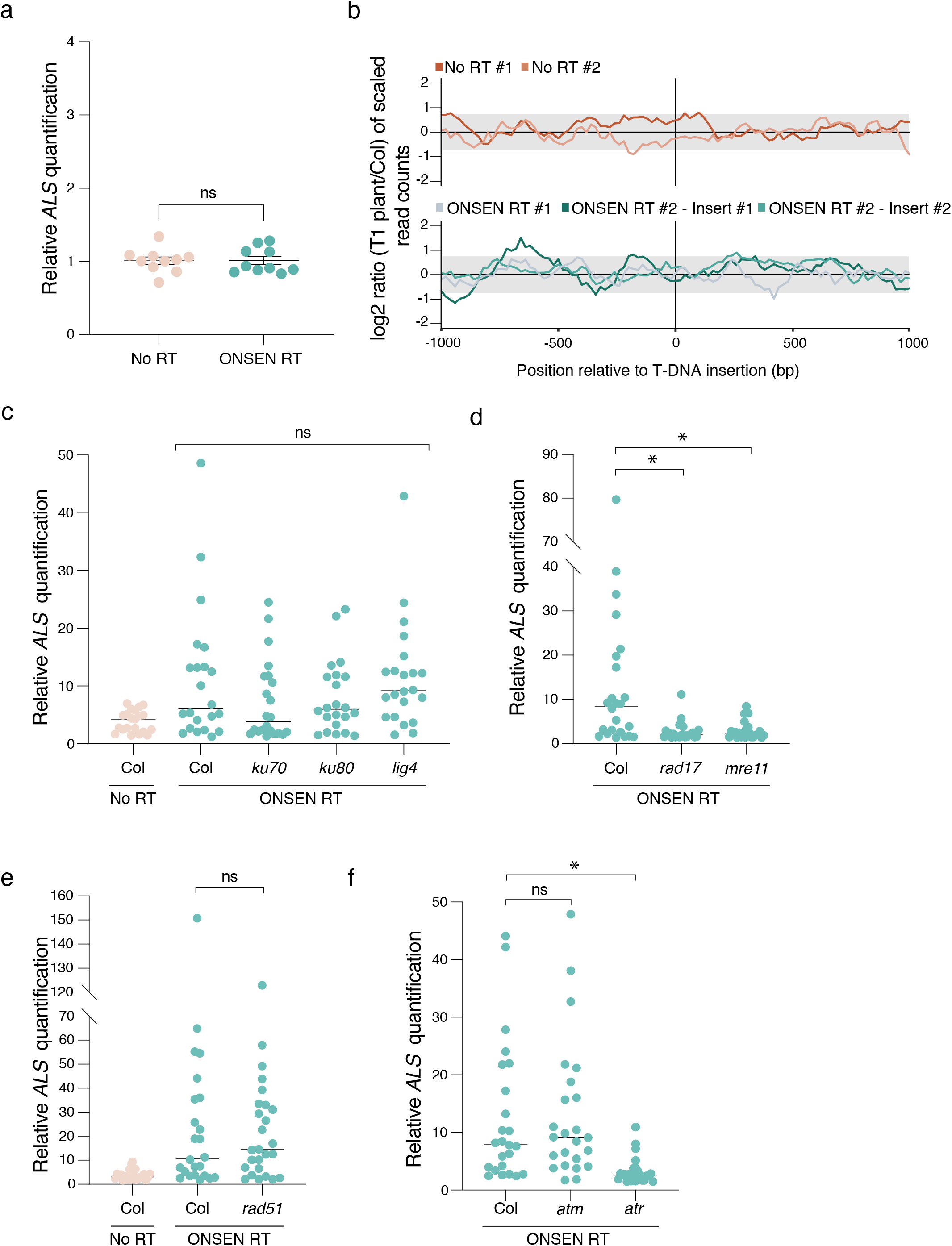
T-DNA amplification is dependent on DNA repair. **a.** DNA-qPCR of *ALS* in No RT- and ONSEN RT-transformed Agrobacterium. Each dot represents a liquid culture initiated from an independent colony. ns = not significantly different (Mann-Whitney *U* test). **b.** log2 ratio between the number of reads corresponding to the genomic regions surrounding the T-DNA insertion sites compared to Col in 20bp bins. The grey box represents the interval where 95% of copy number values from the Arabidopsis genome are present. **c-f.** DNA-qPCR of *ALS* in mutant backgrounds of the (c) NHEJ, (d) TMEJ, and (e) HR pathways, and of (f) DNA damage-induced kinases. Each dot represents an individual T1 plant. Horizontal bars indicate the median. \**P* < 0.01, ns = not significantly different (Kruskal-Wallis ANOVA followed by Dunn’s test).

RT-mediated T-DNA amplification allows us to increase the number of T-DNA copies in Arabidopsis transformants. Higher copy number of T-DNA has the potential to impact many biotechnological applications in plants. To provide a proof-of-concept of the utility of inducing T-DNA amplification, we measured the efficiency of targeted mutagenesis by CRISPR/Cas9 using RT-derived plasmids. We designed and tested three different sgRNAs that target *CRYPTOCHROME 2 (CRY2)* (Extended Data Fig. 6a). Our data indicates that all three sgRNAs induced higher mutation rates, on average, when present on plasmids containing RT sequences (Fig. 4a and Extended Data Fig. 6b-c). In addition, individuals with CRISPR/Cas9-mediated indels displayed higher levels of T-DNA copies than plants with no detectable indels (Extended Data Fig. 6d). Taken together, these results demonstrate the benefits of inducing T-DNA amplification in Arabidopsis to increase targeted mutagenesis rates.

**Figure 4.**
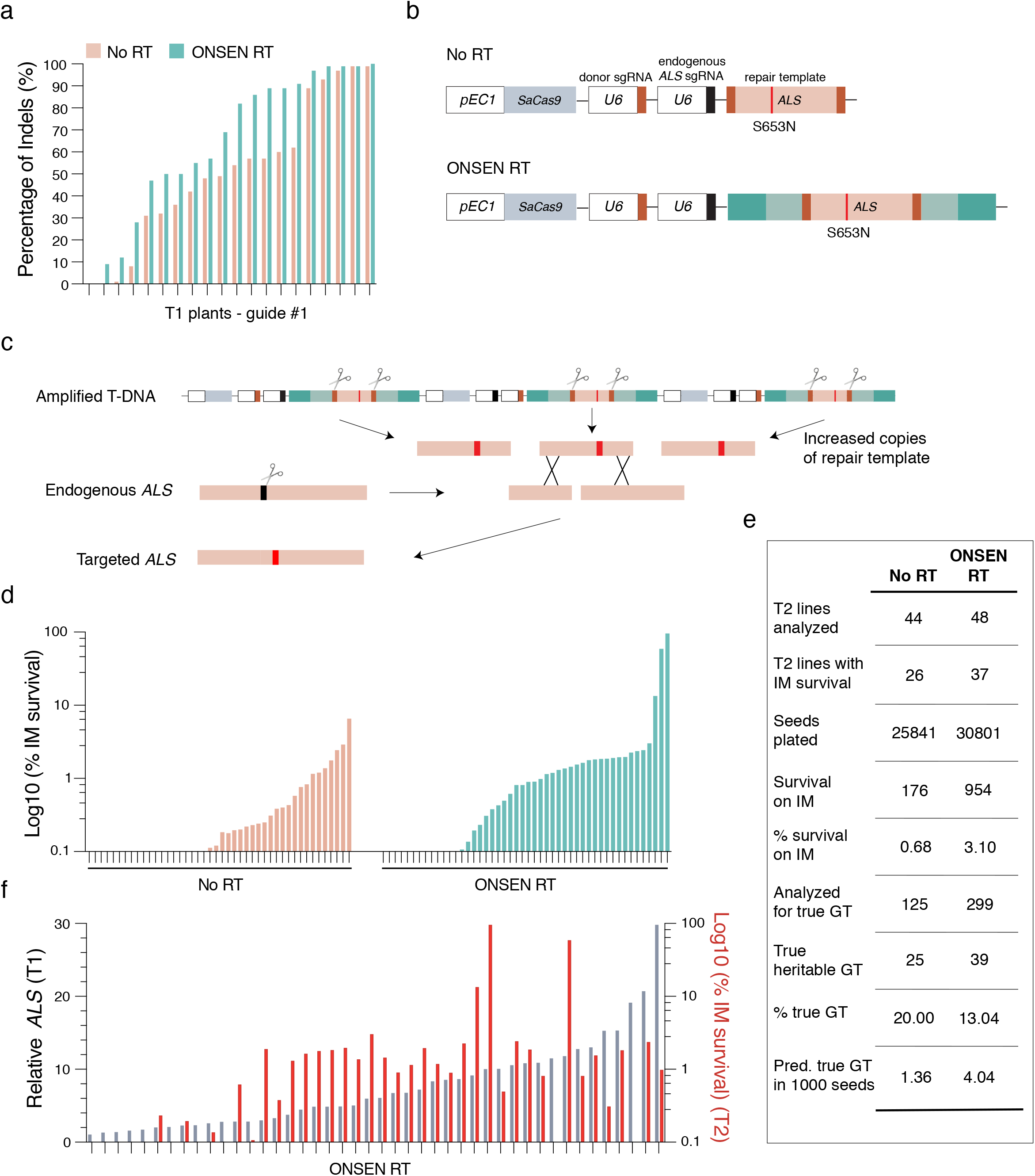
T-DNA amplification increases gene editing efficiency. **a.** Percentage of indels in the *CRY2* gene of DNA extracted from leaves of individual T1 plants transformed with No RT and ONSEN RT constructs containing *CRY2* sgRNA #1. **b.** Schematic representations of constructs used for gene targeting of *ALS*. SaCas9: *Staphylococcus aureus Cas*9. **c.** IPGT approach with ONSEN RT-induced T-DNA amplification. **d.** Percentage of IM-resistant T2 plants from individual T1 parents transformed with No RT or ONSEN RT constructs. **e.** Summary of gene targeting events using No RT and ONSEN RT constructs. **f.** DNA-qPCR of *ALS* in ONSEN RT T1 plants (gray) in relation to percentage of IM-resistance for associated T2 plants (red).

T-DNA amplification may also improve methods used to perform gene targeting in plants. For example, *in planta* gene targeting (IPGT) relies on the chromosomal integration of a T-DNA containing *Cas9*, two sgRNAs genes, and a repair template (Fig. 4b)^25^. One sgRNA is used to create a double-stranded DNA break to initiate DNA repair at a target locus, while the other directs Cas9 to the T-DNA to excise the repair template, which facilitates homology-directed repair (HDR). For single-copy T-DNA integrations, IPGT must rely on only one repair template copy per diploid cell in T1 plants. By inducing T-DNA amplification, more copies of the repair template can be made available for HDR in each cell, which may result in higher gene targeting rates (Fig. 4c). To test this hypothesis, we modified a previously described system targeting the endogenous *ALS* locus of Arabidopsis^11^. Mutating serine 653 to asparagine (S653N) in *ALS* confers resistance to the herbicide imazapyr (IM), thus providing a visual assay to detect and quantify gene targeting events^11^. We designed an IPGT plasmid based on the ALS-IM system that contained the ONSEN RT sequences. Col plants were transformed with the RT-based IPGT plasmid (ONSEN RT) or a standard IPGT plasmid (No RT) (Fig. 4b), and gene targeting rates were measured in T2 seed populations (from individual T1 parents) grown on IM-containing plates. Our results show that more T1 lines produced IM-resistant seedlings when transformed with the RT plasmid (37/48 or 77.1%) compared to the control plasmid (26/44 or 59.1%) (Fig. 4d-e), and that, in general, higher T-DNA copy numbers in T1 plants produced more IM-resistant T2 seedlings (Fig 4e-f). Comparing all T2 seedlings analyzed, we detected a higher percentage of IM-resistant seedlings when they were transformed with the RT plasmid (3.10% versus 0.68%) (Fig. 4e). In these experiments, plants can gain resistance to IM via gene targeting, or through a process known as ectopic gene targeting (EGT). EGT occurs when part of the *ALS* genomic sequence is copied onto the T-DNA to generate a functional *ALS* S653N gene, which is subsequently integrated randomly into the genome^11^. Using the ALS-IM system, EGT can easily be differentiated from true gene targeting events by PCR^11^, and our results indicate that the frequency of true gene targeting events is approximately three times higher when the ONSEN RT plasmid is used (Fig. 4e, Extended Data Fig. 6e). Overall, these results indicate that T-DNA-dependent gene targeting systems in plants can be improved by inducing T-DNA amplification.

In conclusion, this work reveals novel mechanisms that regulate T-DNA structure in Arabidopsis. The ability to regulate T-DNA copy number should allow plant biologists to better control expression levels of transgenes delivered from Agrobacterium, which are subject to RNAi-mediated silencing in plants^3^. It is interesting to note that many binary vectors used for Agrobacterium-mediated transformation contain DNA repeats. For example, the 35S promoter, commonly used to drive constitutive expression of a gene-of-interest or a selection marker in T- DNA, frequently includes two tandem repeated copies of an enhancer sequence^26^. Therefore, re- designing standard binary vectors to avoid DNA repeats may generate more phenotypic consistency among individual T1 lines generated from Agrobacterium-mediated transformation. Conversely, including RT or other DNA repeats on T-DNA provides an opportunity to increase the amount of exogenous DNA integrated at a single locus in plant genomes, which can contribute to useful applications in plant biology as we have shown in this study with gene editing.

## Acknowledgments

We thank all current and former members of the Jacob lab, especially Franziska Langhammer and Gonzalo Villarino, for reagents, discussions, and advice, and Christopher Bolick and his staff at Yale for help with plant growth and maintenance. This project was supported by grant #R35GM128661 from the National Institutes of Health (Y.J.). S.W. was supported in part by an NIH Director’s New Innovator Award (DP2GM137414). We want to thank Dr. Holger Puchta from the Karlsruhe Institute of Technology for his generous gifts of the gene targeting binary plasmids used in this work.

## Contributions

Y.J. supervised the study and designed the experiments with L.D. and W.Y. All experiments were performed by L.D. and W.Y., except the targeted mutagenesis work (C.L.) and the bioinformatic analyses (G.T.). S.W. designed the probes for the FISH experiments. Y.J. and C.L. wrote the manuscript, with contributions from L.D. and W.Y.

## Material and Methods

### Plant materials

Arabidopsis plants were grown under cool-white fluorescent lights (∼100 μmol m^−2^ s^−1^) in long- day conditions (16 h light/8 h dark, 22°C). The T-DNA insertion mutants *atxr5/6* (*At5g09790* / *At5g24330*, SALK_130607 / SAIL_240_H01^20^), *tsk/bru1-4* (*At3g18730*, SALK_034207^27^), *rad51* (*At5G20850*, GK_134A01^28^), *ku70-2* (*At1g16970*, SALK_123114c^29^), *ku80-7* (*At1g48050*, SALK_112921^29^), *lig4-4* (*At5g57160*, SALK_044027^30^), *rad17-2* (*At5g66130*, SALK_009384^31^), *mre11* (*At5g54260,* SALK_028450^32^), *atm (At3g48190,* SALK_040423C^33^), and *atr* (*At5g40820,* SALK_032841C^34^) are in the Col-0 genetic background. They were obtained from the Arabidopsis Biological Resource Center (Columbus, Ohio).

### Cloning

All plasmids used in this study are derived from the DSB/DSB PcUbi4-2 and DSB/DSB AtEC1.1/1.2 vectors previously described^11^. These vectors encode a *Staphylococcus aureus* CRISPR/Cas9 system. The derivative plasmids lacking the ubiquitin promoter and *Cas9* sequence were made using the DSB/DSB PcUbi4-2 plasmid, which was digested using AscI and EcoRI, blunted with T4 DNA Polymerase (New England Biolabs, Ipswitch, MA), and re-ligated using T4 DNA Ligase (NEB). All RT-based plasmids (lacking *Cas9*) were generated by inserting an RT- *ALS* cassette (described below) in place of the *ALS* only cassette using the AatII and PacI restriction sites.

To make the ONSEN RT cassette, ONSEN (*At1g11265*, 4956 bp) was amplified from Col (-109 bp from the beginning of 5’LTR, to +114 bp from the end of the 3’LTR) and cloned into pCR2.1- TOPO (Invitrogen, Waltham, MA). An AscI site was then created in ONSEN at nucleotide position 987-994 (relative to 5’ LTR), replacing GTCACCGT with GGCGCGCC. The ALS repair template and the surrounding sgRNA binding sites were amplified from DSB/DSB AtEC1.1/1.2 and inserted into the pCR2.1-TOPO-ONSEN vector using the AscI and BsrGI sites to generate pCR2.1-TOPO-ONSEN-ALS. The resulting ONSEN-*ALS* cassette was transferred to the binary plasmid (lacking *Cas9*) using the AatII and PacI restriction sites. For the binary plasmids expressing the other RTs, the LTRs and linker sequences of Copia13 (*At2g13940*), Copia21 (*At5g44925*), EVD (*At5g17125*) and GP3-1 *(At3g11970*) were mapped in the Arabidopsis genome and then synthesized at GenScript (Piscataway, NJ). The length of the 5’ linker (546 bp) and 3’ linker sequences (734 bp) synthesized for each RT was based on the length of the linker sequences in the ONSEN RT vector. A multicloning sequence that includes a NotI restriction site was inserted between the 5’ and 3’ linker sequences of each RT. The ALS repair template in DSB/DSB AtEC1.1/1.2 was removed from the vector using NotI and inserted in the NotI cloning site of the four synthesized RTs. Finally, the RT-*ALS* cassettes were transferred to the binary plasmid (lacking *Cas9*) using AatII and PacI.

The series of truncated ONSEN RT plasmids were created from a synthesized (GenScript) ONSEN-*ALS* cassette cloned into pUC57. The sequence of this synthesized ONSEN-*ALS* cassette is identical to the ONSEN-*ALS* cassette described in the previous paragraph, except for the insertion of restriction sites (each one cutting only at a single location) right before and after the different sections of ONSEN-*ALS*: 5’ LTR, 5’ linker, ALS, 3’ linker, and 3’ LTR. To remove a specific section of the synthesized ONSEN-*ALS* cassette, two restriction enzymes targeting the borders of that section were used to digest pUC57-ONSEN-*ALS*. The digested plasmid was then blunted using Quick Blunting kit (NEB) and re-ligated using T4 DNA Ligase (NEB) to generate the deletion. The random repeat sequence was generated using the online tool Random DNA Sequence Generator online tool (The Maduro Lab, UC Riverside), synthesized at GenScript, and cloned into pUC57-ONSEN-*ALS*, replacing the 5’ and 3’ LTRs. Finally, all modified ONSEN- *ALS* cassettes were cloned into the binary plasmid (lacking *Cas9*) using AatII and PacI. The binary plasmid lacking one of the two sgRNA genes was generated by digesting the No RT plasmid with XmaI and PacI, blunting (Quick Blunting kit, NEB), and re-ligating using T4 DNA Ligase (NEB).

The binary plasmids (No RT and ONSEN RT) used for transformation into the Arabidopsis mutant backgrounds were modified to replace the kanamycin resistance gene (*NPTII*) with an hygromycin resistance gene. Briefly, the *NPTII* gene cassette was removed from the No RT and ONSEN RT plasmids using the restriction enzymes AatII and PmeI. In parallel, the *NPTII* gene cassette (including promoter and terminator) was cloned into pAGM1311 and modified using HiFi DNA Assembly Cloning Kit (NEB, Ipswitch, MA) to replace the complete coding sequence of *NPTII* with the coding sequence of the hygromycin resistance gene from pMDC7^35^. The modified cassette was then reinserted into the No RT and ONSEN RT plasmids.

For the gene editing experiments involving the detection of mutations at the *CRY2* locus, the DSB/DSB PcUbi4-2 plasmid was first modified by replacing the original ALS cassette with an ALS cassette lacking the sgRNA binding sites using AatII and PacI. The sgRNA binding sites were eliminated to prevent cutting of the T-DNA locus by the Cas9-sgRNA complex, which could affect T-DNA quantification. To build the equivalent ONSEN RT plasmid, an ALS cassette without the sgRNA binding sites was inserted in the pCR2.1-TOPO-ONSEN vector using the AscI and BsrGI sites, and the resulting ONSEN-ALS cassette was then cloned into the DSB/DSB PcUbi4-2 plasmid at the AatII and PacI sites. Finally, the sgRNA gene targeting the *ALS* endogenous locus was replaced by a sgRNA gene targeting *CRY2*. This was done by digesting the ONSEN RT plasmid with XmaI and PacI and subcloning the fragment (containing the endogenous *ALS* gRNA gene) into a pENTR/D-Topo vector (Thermo fisher Scientific, Waltham, MA). The resulting plasmid was PCR amplified with a primer pair to change the *ALS* sgRNA spacer sequence to one of three different *CRY2* sequences (sgRNA#1, 5’-AAGATCGCTGAAATCGTGTT-3’; sgRNA#2, 5’-GCAGGACCGGTTATCCGTTG-3’; and sgRNA#3, 5’- CCGATCATGATCTGTGCTTC-3’). The amplified PCR products were then ligated with T4 DNA Ligase (NEB) and the *CRY2* sgRNA genes were subcloned into the modified DSB/DSB PcUbi4-2 plasmids (i.e., lacking sgRNA binding sites) with XmaI and PacI.

To produce the ONSEN RT plasmid for gene targeting, the ONSEN RT cassette (containing ALS and the sgRNA binding sites) was amplified from the pCR2.1-TOPO-ONSEN-ALS vector and inserted into DSB/DSB AtEC1.1/1.2 using the AatII and PacI sites, replacing the ALS only cassette. The original DSB/DSB AtEC1.1/1.2 plasmid (No RT plasmid) served as a control.

### Plant transformation

Arabidopsis plants were transformed by using the floral dip method^36^. Briefly, one day prior to floral dip transformation, 300 μL of a stationary Agrobacterium (strain GV310) liquid culture was used to inoculate 200 mL of LB containing 100 mg/L gentamycin, 100 mg/L spectinomycin. The culture was incubated with shaking overnight at 28°C. The bacterial culture was spun down at 3,220 x g for 25 min and resuspended in 200 mL of transformation solution (5% sucrose and 0.02% Silwet L-77). Arabidopsis flowers were dipped into the bacterial solution, gently agitated for 10 seconds, then stored horizontally in a tray with a blackout lid overnight in a long-day growth chamber. T1 plants were selected on ½ MS plates containing 1% sucrose, carbenicillin (200 μg/mL) and either kanamycin (100 μg/mL) or hygromycin (25 μg/mL). Herbicide-resistant seedlings were transferred to soil after 7-10 days on plates. Transformed DNA repair mutants were all homozygous except for *rad51*, due to sterility^28^. We therefore transformed *rad51* heterozygous plants and analyzed homozygous T1 mutants.

### Plant DNA extraction

Leaves from plants that were grown for 2 weeks on soil (unless otherwise indicated) were homogenized in 500 μL DNA extraction buffer (200 mM Tris-HCl pH 8.0, 250 mM NaCl, 25 mM EDTA, and 1% SDS) and 50 μL phenol:chloroform:isoamyl alcohol (25:24:1). Each sample was centrifuged for 7 min at 16,000 x g, and 300 μL of the aqueous layer was transferred to a 1.5 mL tube containing 300 μL isopropanol. Samples were vortexed, incubated at room temperature for 5 min, and then spun down at 16,000 x g for 10 min. The supernatant was removed and the pellets were washed with 400 μL 70% ethanol. After centrifugation at 16,000 x g for 5 min, the ethanol was removed and the pellets were dissolved in 100 μL of water.

### Agrobacterium DNA extraction

GV310-transformed colonies were grown in liquid culture overnight at 28°C. 300 μL of each culture was transferred to a 1.5 ml tube and centrifuged for 1 min at 16,000 x g. The resulting pellet was resuspended in 250 μL of a lysozyme solution (200 mM CaCl_2_ with 1% lysozyme). The samples were incubated at 42°C for 5 min and 750 μL of 96% ethanol was added, followed by centrifugation for 10 min at 16,000 x g. The supernatant was removed, and the pellet was air dried for 10 min and resuspended in 100 μL of water.

### DNA-qPCR

Real-time PCR was carried out using a CFX96 Real-Time PCR Detection System (Bio-Rad, Hercules, CA) with a KAPA SYBR FAST qPCR Master Mix (2×) Kit (Kapa Biosystems, Wilmington, MA). Relative quantities were determined by the 2(-delta delta C_t_) method^37^ using *Actin7* (*At5g09810*) and the gene coding for hypothetical protein WP_046033610.1 (NCBI accession number) as the normalizers for DNA extracted from Arabidopsis and Agrobacterium, respectively. Relative quantities of *ALS* in Arabidopsis plants were calculated using Col DNA as the calibrator. For relative quantification of *pea3A(T)* and the *NPTII* cassette in Arabidopsis and of *ALS* in Agrobacterium, a DNA sample from a “No RT” transformant was used as the calibrator.

### Fluorescence *in situ* probe design

Primary FISH probes were designed with a procedure previously described^38^, with some modifications. First, a pool of primary targeting sequences were designed with OligoArray2.1^39^, with the following parameters: sequence length 30 nt; minimum melting temperature 66°C; maximum melting temperature 100°C; secondary structure melting temperature limit 76°C; cross-hybridization melting temperature limit 72°C; minimum GC content 30%; maximum GC content 90%; avoiding 6 or more consecutive A, T, G or C’s; allowing at most 20-nt overlap between adjacent target sequences. The primary targeting sequences were compared against the TAIR10 genome to ensure specificity. Then, a 30 nt secondary probe binding sequence (reverse complement of the dye-labeled secondary probe sequence) was appended to the 3’ end of each primary targeting sequence to generate the full-length primary probes (Integrated DNA Technologies, Coralville, IA). To detect the *ALS* and the *NPTII* genes, 39 and 29 primary probes were designed, respectively (Supplementary Table 1). The secondary probe sequences were adapted from a previous report^38^. The secondary probes for *ALS* and *NPTII* were conjugated to 5’ Alex Fluor 488 and ATTO 590, respectively (IDT).

### Fluorescence *in situ* hybridization

Leaves from 3-week-old plants were fixed in cold 4% formaldehyde in Tris buffer (10 mM Tris-HCl pH 7.5, 10 mM NaEDTA, 100 mM NaCl) for 20 min, and then washed twice in Tris buffer. The leaves were chopped with a razor blade in 500 μl LB01 buffer (15 mM Tris-HCl pH7.5, 2 mM NaEDTA, 0.5 mM spermine-4HCl, 80 mM KCl, 20 mM NaCl and 0.1% Triton X-100), and the resulting slurry was filtered through a 30 μM mesh (Sysmex Partec, Gorlitz, Germany). The filtered solution was mixed 1:1 with sorting buffer (100 mM Tris-HCl pH 7.5, 50 mM KCl, 2mM MgCl2, 0.05% Tween-20 and 5% sucrose), spread onto a coverslip, and dried. Cold methanol was added to the coverslips for 3 min, followed by TBS-Tx (20 mM Tris pH 7.5, 100 mM NaCl, 0.1% Triton X-100). 0.1 mg/ml of RNase A in 2x SSC (0.3M NaCl, 30 nM sodium citrate, pH 7.0) was added onto the coverslips and incubated at 37°C for 45 min, followed by two washes in 2x SSC. Pre-hybridization buffer (2X SSC, 50% Formamide and 0.1% Tween 20) was added for 30 min at room temperature. Probes were added to the hybridization buffer at a concentration of 1 uM, which was applied to each coverslip. The coverslips were then incubated at 80°C for 3 min and at 37°C overnight in a humid chamber. Coverslips were washed twice with SSC-0.1% Tween at 60°C for 15 min, and once for 15 min at room temperature. Secondary probes at a concentration of 40 nM in buffer (2x SSC + 40% formamide) were added to coverslips. The coverslips were incubated at room temperature for 30 min, washed with secondary wash buffer (2x SSC + 40% formamide), and washed twice with 2x SSC. Coverslips were mounted on microscope slides with Vectashield containing DAPI (Vector Laboratories, Burlingame, CA). Nuclei were imaged under a Nikon Eclipse Ni-E microscope with a 100X CFI PlanApo Lamda objective (Nikon, Minato City, Tokyo, Japan) and an Andor Clara camera. Z-series optical sections of each nucleus were obtained at 0.3 μm steps. Images were deconvolved by FIJI using the DeconvolutionLab plugin^40, 41^. The nuclei selected for imaging were :<Σ 55 μm^3^ to enrich for 2C nuclei^42^. The nuclear volume was measured using FIJI with the 3D ImageJ suite^41, 43^.

### Library construction, sequencing and bioinformatic analyses

DNA sequencing libraries were prepared at the Yale Center for Genome Analysis. Genomic DNA was sonicated to a mean fragment size of 350 bp using a Covaris E220 instrument (Covaris, Woburn, MA) and libraries were generated using the xGen Prism library prep kit for NGS (Integrated DNA Technologies, Coralville, IA). Paired-end 150 bp sequencing was performed on an Illumina NovaSeq 6000 using the S4 XP workflow (Illumina, San Diego, CA). Raw FASTQ files were pre-processed and trimmed using the fastp tool^44^ (--length_required 20 --average_qual 20 --detect_adapter_for_pe -w 10). Subsequently, the command line program grep (Free Software Foundation, Boston, MA) was used to search FASTQ files using the 20 bp sequences inside the T-DNA neighboring the LB and RB sites (and their reverse complement) as a query. The filtered sequences were then processed using BioPython^45^ to isolate flanking sequence tags (FSTs) adjacent to the LB and RB. FSTs were then used as BLAST queries^46^ to identify regions of the genome where T-DNA sequences were inserted. Further supporting reads were obtained by mapping the FASTQ files to the genome (TAIR10^47^) using BWA-MEM^48^ and isolating reads for which only a single read of a mate pair maps to the genome at the identified point of insertion, as the other read will map to the T-DNA insertion. Assembling the T-DNA insertion junctions with the genome was then done using the MAFFT multiple sequence aligner^49^, and a reference sequence was manually created. Insert sites were confirmed using PCR and Sanger sequencing.

Estimation of transgene sequence copy number was achieved by mapping reads to a panel of 7,535 genes, along with three regions of the T-DNA transgene. The selected genes are reported in the PLAZA 5.0 database^50^ to originate from single gene families in the genome (omitting plastid genes and *At3g48560* from which the *ALS* sequence of the transgene derives). The read counts from these alignments were extracted using SAMtools idxstats function^51^ and normalized to Bins Per Million mapped reads (BPM; [Reads Per Kilobases]/[sum(Reads Per Kilobases)*1^e6]). The mean BPM of the selected genes were taken as the normalized copy number of a single copy gene in the genome and the relative fold change, in BPM, for regions of the T-DNA transgene was calculated using this as a reference.

Measurement of coverage around mapped T-DNA insertion sites was done using the bamCompare function of the deepTools package^52^ which scales by read count and returns the log2 ratio of two alignments for a genome split into equally sized bins. The bin size was set to 20bp and the genome alignment of each line was compared with a wild-type Col sample sequenced at the same time.

Internal T-DNA junctions were assembled in a similar manner to the identification of insertion junctions with the genome, but using reads with FSTs that match the binary vector used to transform the plants. Once assembled, the FASTQ files were searched again with consensus intersection sequences (+/- 15 bp from either the point of intersection or edge of filler sequence) and only intersections with two or more independent read pairs supporting it were retained.

### Gene editing

To assess the efficiency of targeted mutagenesis of *CRY2*, genomic DNA was extracted from individual 2-week-old T1 plants and used to amplify the *CRY2* gene. The resulting PCR products were sequenced and analyzed for INDEL frequency by Inference of CRISPR Edits (ICE) analysis (Synthego Performance Analysis, ICE Analysis. 2019. V3.0. Synthego).

To measure IPGT rates, T2 seeds from individual T1 lines were plated on ½ MS plates containing 5 µM Imazapyr (IM) and 1% sucrose. Seeds were stratified for 3 days at 5°C. The seed count for each plate was determined by using the ImageJ ‘analyze particles’ function after binary processing. The seeds were allowed to germinate and grow for one week in long day conditions and the imazapyr resistant seedling were then counted manually.

The true gene targeting events were characterized as previously described^11^, with some modifications. Briefly, the endogenous *ALS* locus was amplified with primers specific for regions outside of the repair template and the PCR product was Sanger sequenced. The codon corresponding to amino acid 653 and the gRNA binding site were analyzed for a gene targeting event using Sequencher 5.4.6 (Gene Codes Corporation, Ann Arbor, MI). Samples with different types of editing events were re-analyzed using Synthego ICE or subcloning and further Sanger sequencing. Plants were considered as having undergone true gene targeting when samples displayed, at a minimum, ∼50% GT-edited sequences.

### Data availability

Raw sequencing data are deposited at the NCBI SRA (accession code: PRJNA892619).

### Primers

All primers used in this study are in Supplementary Table 2.

---

**Extended Data Figure 1.**
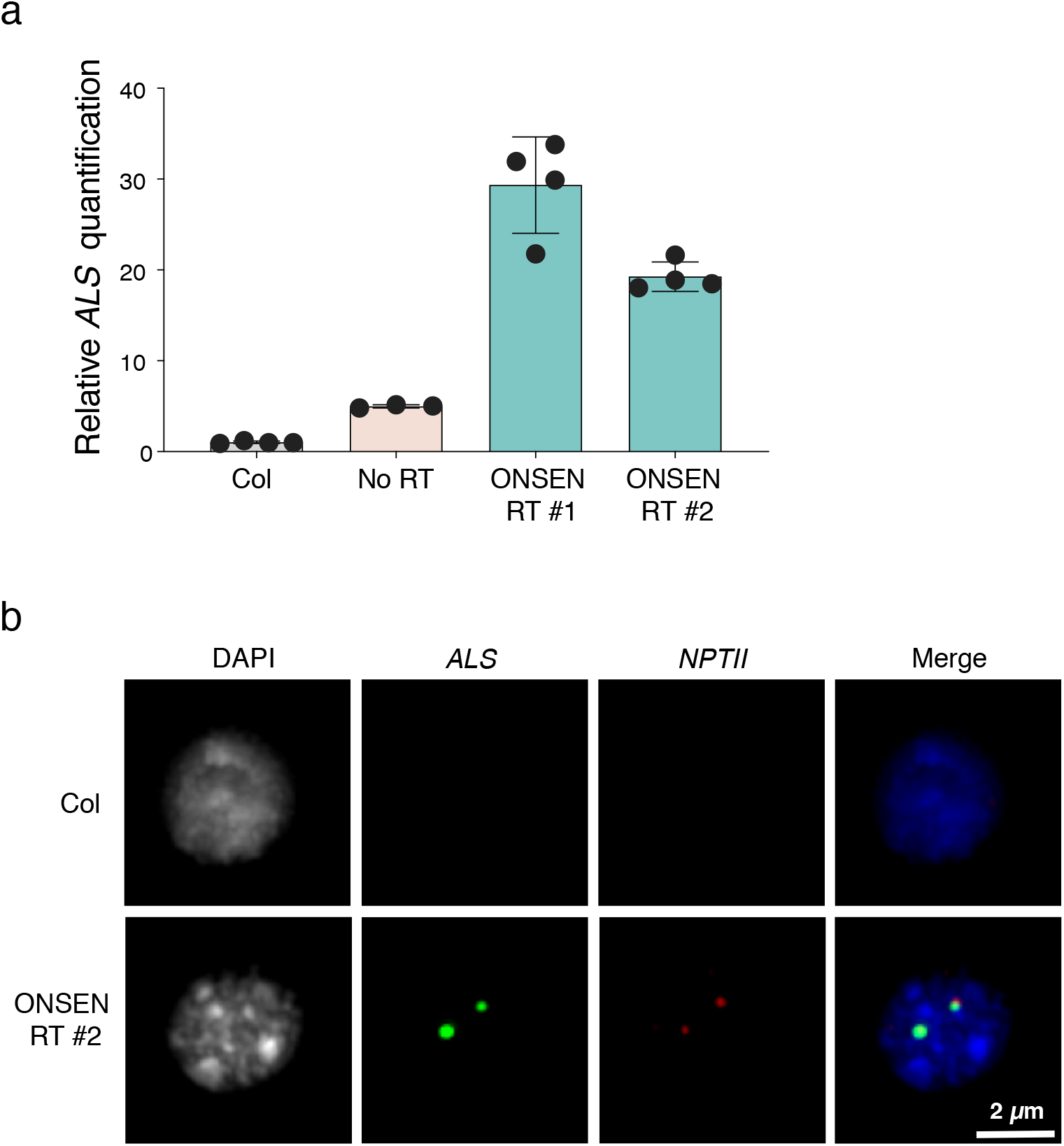
DNA FISH analysis. **a.** DNA q-PCR of *ALS* in Col and T1 plants used for the FISH experiment (panel b and Fig. 1e). Dots represent DNA samples from individual leaves from the same T1 plant. Bars indicate the mean. SD is shown. **b.** FISH in leaf nuclei targeting the T-DNA sequence. Nuclei were stained with DAPI and probes for *ALS* (green) and *NPTII* (red).

**Extended Data Figure 2.**
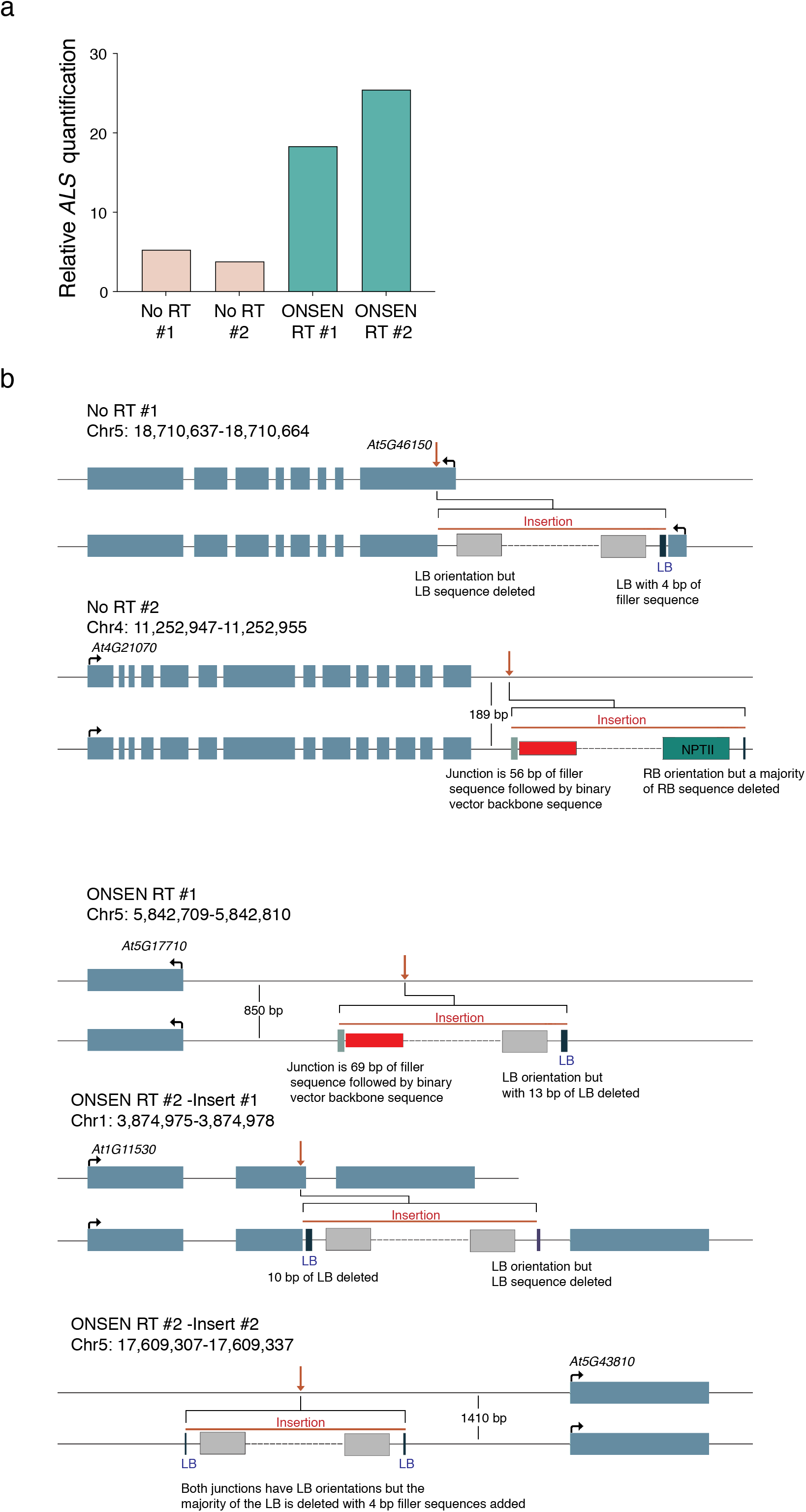
Whole genome sequencing analysis of T-DNA insertions. **a.** DNA-qPCR of *ALS* in the No RT and ONSEN RT T1 plants used for whole genome sequencing (panel b and Fig. 1f-g). **b.** Locus diagrams for the identified T-DNA insertions. The coordinates for each insertion are based on the TAIR10 annotation and correspond to the Arabidopsis genomic borders surrounding each identified T-DNA. For each insertion, top lines represent unaltered genomic sequence with annotated genes. Red arrows represent insertion points. The bottom lines show the borders of the insertion in more detail, with the identified binary vector (non-T-DNA region) or T-DNA components shown. Dashed lines represent contiguous T-DNA-associated cassettes. Red bars indicate binary vector sequence (non-T-DNA), dark blue LB bars to light gray *Pea3A* terminator sequence bars indicate the 5’ end of the T-DNA construct (though it may be in 5’ or 3’ orientation in the plant genome). Darker gray bars adjacent to the red bars are filler sequences, and the teal bar represents *NPTII* sequence.

**Extended Data Figure 3.**
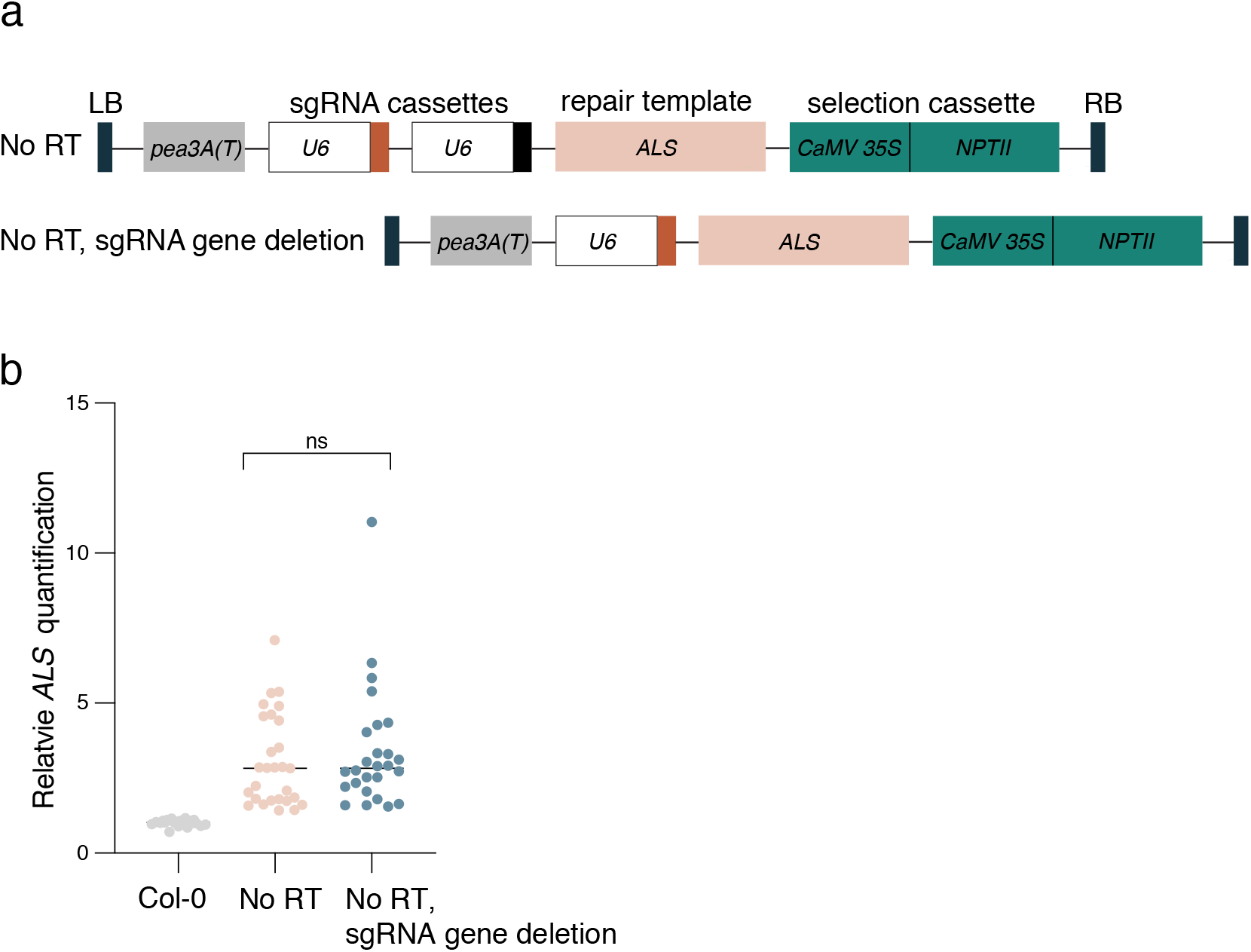
Repetitive sgRNA genes do not contribute to T-DNA amplification. **a.** Schematic representation of the sgRNA gene deletion from the No RT vector. **b.** DNA-qPCR of *ALS* in Col and in T1 plants transformed with the No RT plasmid and the No RT plasmid with a gRNA gene deleted. Each dot represents and individual plant. Horizontal bars indicate the median. ns = not significantly different (Mann-Whitney *U* test).

**Extended Data Figure 4.**
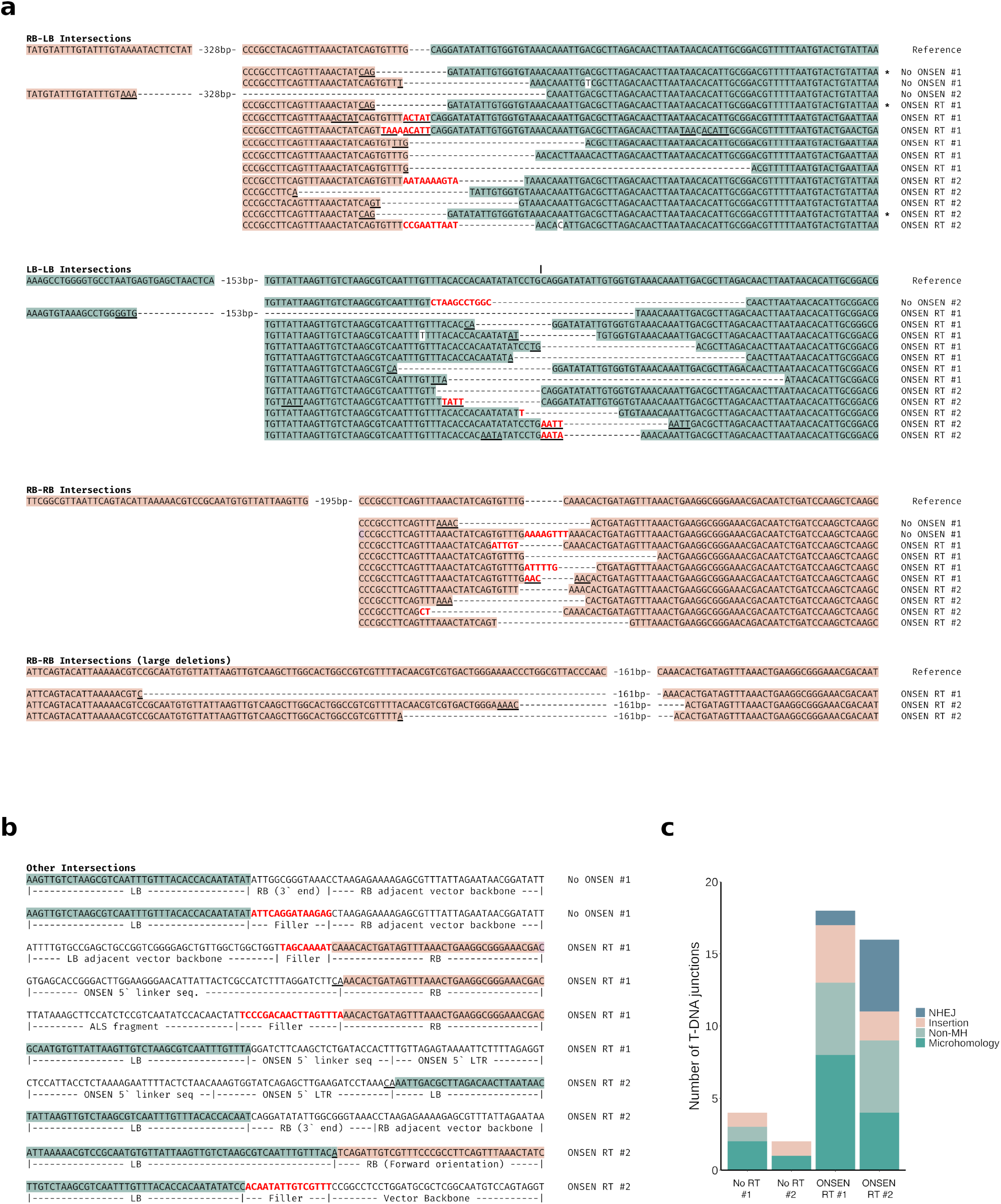
Concatenated T-DNA junctions support the involvement of TMEJ in T-DNA amplification. **a.** Multiple sequence alignments of junctions between two T-DNA sequences grouped by orientation of the two sequences (RB-LB, LB-LB or RB-RB). Reference sequences were constructed assuming T-DNA molecules began and ended immediately upstream of the endonuclease recognition sequence (5′-CAGGATATATT-3′)^53, 54^. Filler sequences are in red and sequences consistent with microhomology-associated deletions are underlined. If filler sequences have a similar sequence nearby, they (and the nearby sequence) are also underlined. Asterisks indicate identical junctions occurring in independent plants. **b.** Depiction of T-DNA junctions with another T-DNA or binary vector sequence not immediately internally adjacent to the LB or RB. **c.** Classification of RB-LB, LB-LB and RB-RB sequences for each T1 line following a procedure previously described^55^. NHEJ (<4 bp deletions and <5bp insertions), insertions (≥5bp with any deletion), Non-Microhomology (Non-MH; ≥4bp deletion or <5bp insertions with microhomologies <2), and Microhomology (MH; ≥4bp deletion with microhomologies ≥2). The latter three are associated with DNA polymerase theta.

**Extended Data Figure 5.**
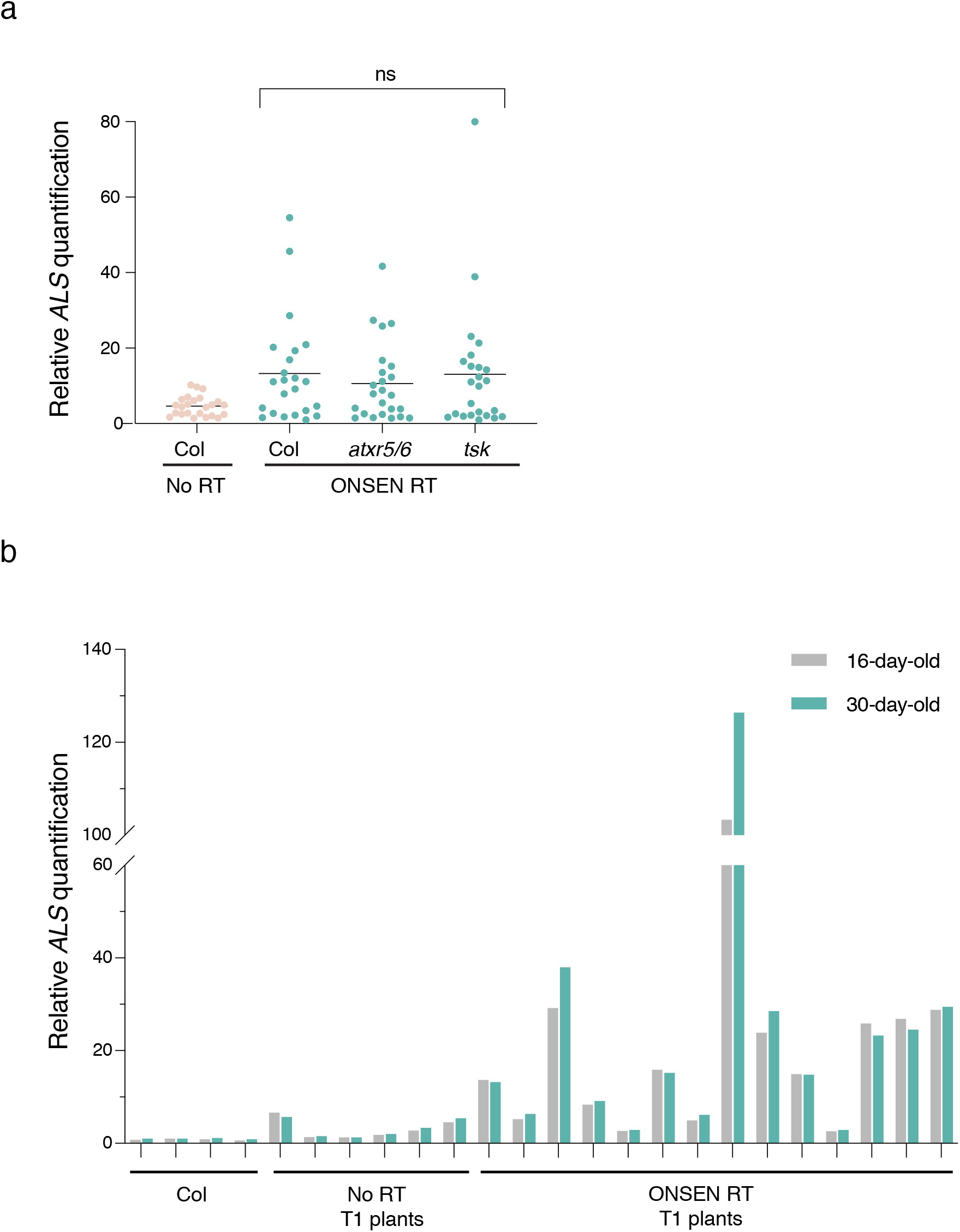
T-DNA amplification is not DNA replication-dependent. **a.** DNA-qPCR of *ALS* in Col, *atxr5/6,* and *tsk* plants transformed with the ONSEN RT construct. Each dot represents an individual T1 plant. Horizontal bars indicate the median. ns = not significantly different (Kruskal-Wallis ANOVA followed by Dunn’s test). **b.** DNA-qPCR of *ALS* in DNA extracted from leaves of T1 plants at 16 days and 30 days after germination.

**Extended Data Figure 6.**
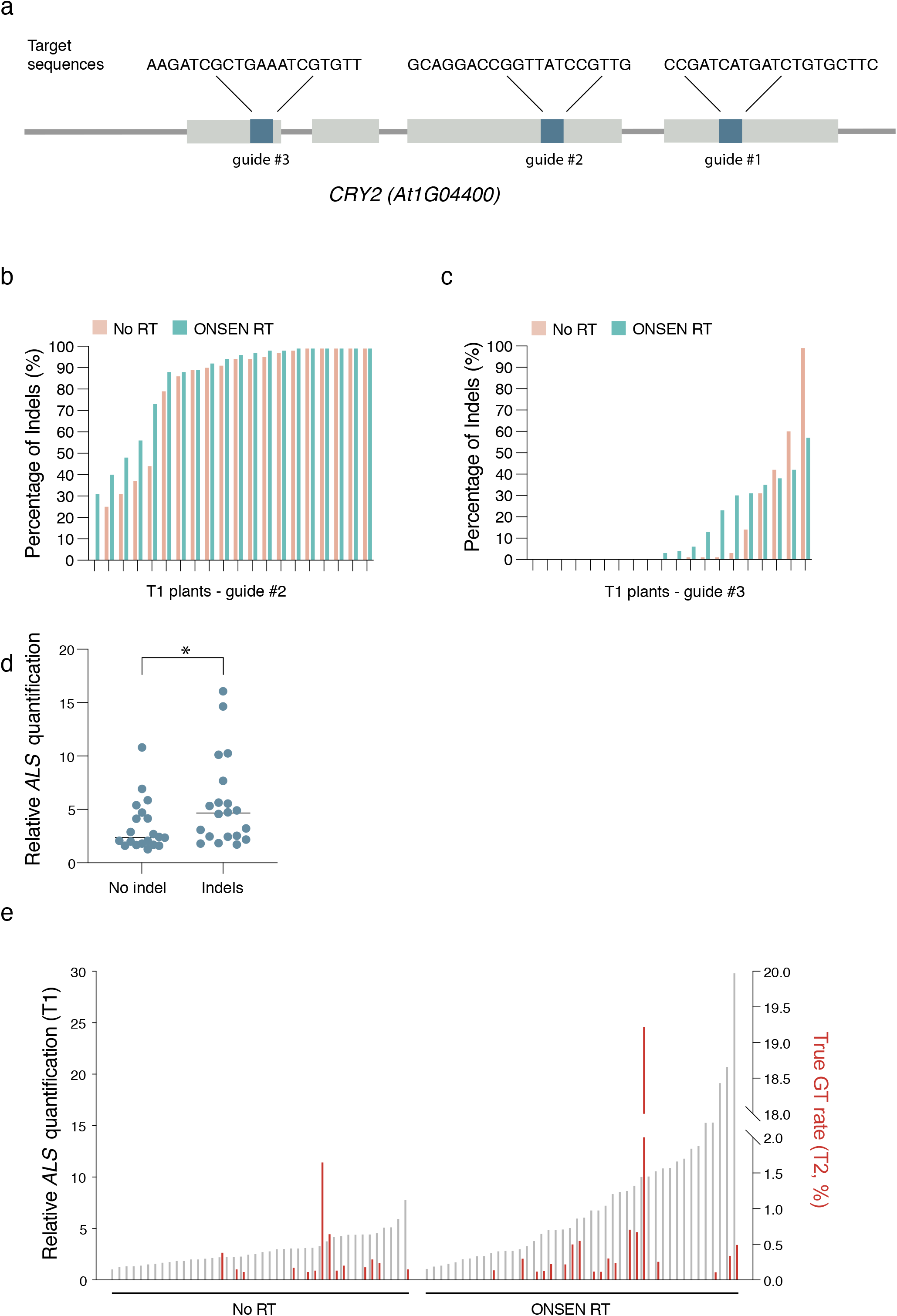
T-DNA amplification increases the efficiency of targeted mutagenesis and gene targeting. **a.** Gene structure of *CRY2.* Gray bars represent exons, and blue bars represent regions targeted by sgRNAs. **b-c.** Percentage of CRISPR-induced indels in leaf tissue of individual T1 lines transformed with either the No RT or the ONSEN RT construct. Constructs carried either (b) sgRNA #2 or (c) sgRNA #3. **d.** DNA-qPCR of *ALS* in T1 plants with no detectable *CRY2* indels (No indels) or detectable indels (Indels) using CRY2 guide #3. **e.** DNA-qPCR of *ALS* in No-RT and ONSEN-RT T1 plants (gray) in relation to the percentage of true (non-ectopic) gene targeting rates in the T2 generation (red).

